# Litter expansion alters metabolic homeostasis in a sexually divergent manner

**DOI:** 10.1101/2020.07.23.217307

**Authors:** Kavitha Kurup, Shivani N Mann, Jordan Jackson, Stephanie Matyi, Michelle Ranjo-Bishop, Willard M. Freeman, Michael B Stout, Arlan Richardson, Unnikrishnan Archana

## Abstract

Nutritional manipulations early in life have been shown to influence growth rate and elicit long lasting effects which in turn has been found to impact lifespan. Therefore, we studied the long-term effects of pre-weaning dietary restriction implemented by litter expansion (4, 6, 8, 10, and 12 pups per dam: LS4, LS6, LS8, LS10, LS12) on male and female C57BL/6 mice. After weaning, these mice were fed *ad libitum* a commercial lab chow for the 15-month duration of the study. The mice from large litter sizes (LS12) were significantly leaner and had reduced total fat mass compared to the normal size litters (LS 6) starting from weaning through to 15 months of age. Male LS10 & 12 mice showed significant reduction in their fat depot masses at 15 months of age: gonadal, subcutaneous, and brown fat whereas the females did not mimic these findings. At 9 months of age, both male and female LS10 and 12 mice showed improved glucose tolerance; however, only male LS10 and LS12 mice showed improved insulin tolerance starting at 5 months of age. In addition, we found that the male LS8, 10 & 12 mice at 15 months of age showed significantly reduced IGF-1 levels in the serum and various other organs (liver, gastrocnemius and brain cortex). Interestingly, the female LS8, 10, 12 mice showed a different pattern with reduced IGF-1 levels in serum, liver and gastrocnemius but not in the brain cortex. Similarly, the litter expanded mice showed sexual divergence in levels of FGF21 and adiponectin with only the male mice showing increased FGF21 and adiponectin levels at 15 months of age. In summary, our data show that, litter expansion results in long-lasting metabolic changes that are age and sex dependent with the male mice showing an early and robust response compared to female mice.

## Introduction

Nutrition has a major environmental influence during pre-natal and post-natal development that can lead to major long-term effects both physically and mentally later in life. Barker hypothesized that intrauterine growth retardation increases the risk to many chronic diseases such as diabetes, metabolic syndrome and cardiovascular disease in middle to later life [1, 2, 3]. Maternal under-nutrition during pregnancy permanently alters the fetal organ structure leading to developmental programming wherein the fetal metabolism changes to ensure survival and results in intrauterine growth restriction and low-birth weight [4]. Offspring born with low-birth weights have been shown to develop obesity, insulin resistance, type 2 diabetes and coronary artery disease during adulthood in both humans and animal models [5, 6, 7, 8]. Ozanne and Hanes [9] showed that poor maternal nutrition during pregnancy (reduced protein to 8%) in mice results in low-birth weight pups and when these low-birth weight pups were cross fostered to dams fed a normal diet (20% protein) they showed reduced longevity (∼25%), and when these pups were fed western diet after weaning, it further shortened their lifespan. In the opposite scenario, when mothers are over-nourished during their pregnancy phase (leading to maternal obesity and gestational diabetes), the offspring are at increased risk to develop obesity, metabolic syndrome and cardio vascular disease later in life [4, 10, 11, 12, 13].

In contrast to the above studies, Ozanne and Hales [9] showed that pups born to mothers on normal diet (20% protein) when cross fostered to lactating mothers on low-protein (8%) diet (exposure to restricted diet only during lactation) and then fed a normal chow ad libitum after weaning, show slow post-natal growth and have increased longevity (6 %). Furthermore, when these pups were exposed to western diet after weaning, they were resistant to reductions in lifespan when compared to pups born to mothers with low protein diet [9]. Additionally, Lopez-Soldadao et al. [14] reported that litter expansion during lactation phase (without any nutritional manipulation to the dams) created dietary restriction in the pups by showing that rat pups from large litters consumed less milk than the controls and that the pups from large litters showed reduced adiposity and improved insulin sensitivity during adult life. More recently, Sun et al. [15] studied the effect of dietary restriction during the pre-weaning period (3 weeks of lactation) on lifespan. The authors implemented dietary restriction during lactation in two ways: (i) lactating mothers were fed low protein diet similar to what Ozanne and Hales [9] had done previously and (ii) by litter expansion, i.e., increasing the litter size by 50% (8 vs 12 pups/litter) on lactating dams on normal, high protein (20%) diet. They found that maternal protein restriction had no effect on the lifespan of the offspring; however, the litter expansion by 50% resulted in a significant (∼18%) increase in lifespan of the pups subjected to litter expansion. Subsequently, Sadagurski et al. [16] showed that litter expansion in mice had long-lasting beneficial effects such as improved energy homeostasis and insulin and leptin sensitivity throughout the lifespan of the mice. Cumulatively, these studies clearly show that under or over-nutrition during intrauterine growth results in reduced lifespan and increased risk to various chronic diseases whereas reduced nutrition only during the post-natal phase leads to slow post-natal growth and increased lifespan.

The purpose of this study was to comprehensively evaluate the long-term effects of litter expansion. We evaluated the long-term (15 months) effects of a variety of litter sizes (LS4, LS6, LS8, LS10 and LS12) on metabolic parameters in both male and female C57BL/6J mice. Along with many metabolic indices including body composition, adiposity and insulin sensitivity we also assessed the levels of growth factors (IGF-1 & FGF21) and adiponectin. Interestingly, all long-lasting beneficial effects observed with litter expansion were found to be sexually divergent. Male mice displayed early and robust response, whereas female mice demonstrated a delayed but significant response later in life.

## Materials and Methods

### Animals

Litter expansion of male and female C57BL/6J mice was performed by Jackson laboratory (Bar Harbor, ME) and 5 litter sizes (LS) were generated: 4, 6, 8, 10, and 12 pups per litter (LS4, LS6, LS8, LS10, and LS12). In brief, 6 pups/litter was used as control as that is the average litter size of C57BL/6J mice at Jackson Laboratory. To generate the other litter sizes, pups were added to lactating dams to generate litter sizes 8, 10, and 12 pups/litter and reduced to generate 4 pups/litter. No rejection of pups by the dams were noticed in the larger litter sizes. All lactating mothers were fed the standard breeding chow rich in protein (∼18%) and fat (10-12%). At 3 weeks of age, all the male and female mice were shipped to the University of Oklahoma Health Sciences Center and maintained under SPF conditions in a HEPA barrier environment. The animals were maintained (5 mice/cage) under temperature and light controlled conditions (12-12h light-dark cycle) and fed *ad-libitum* irradiated NIH-31 mouse/rat diet from Teklad (Envigo, Madison, WI). Each group contained 10 to 12 animals and these mice were separated into two cohorts, one for body weight and insulin sensitivity and one cohort for rest of the molecular analysis (Gene expression, IGF-1, FGF21 and adiponectin levels). For the molecular analysis mice were taken at 15 months of age, fasted overnight, sacrificed and tissues harvested, snap frozen in liquid nitrogen, and stored at -80°C until used. All procedures mice were approved by the Institutional Animal Care and Use Committee at the University of Oklahoma Health Sciences Center.

### Sexual maturity

Female sexual maturity was determined by following the onset of vaginal patency [17]. The mice were observed for the visual appearance of an opening starting at postnatal day 21 to postnatal day 32. The appearance of opening was defined as the onset of vaginal opening.

### Body Composition

Body composition of the crowded litter animals was measured using nuclear magnetic resonance spectroscopy (NMR-Bruker minispec) at 15 months of age. Fat mass and lean body mass were measured.

### Liver Triglyceride Content

Liver samples (∼100 mg) were homogenized on ice for 60 seconds (20 second increments) in 10X (v/w) Cell Signaling Lysis Buffer (Cell Signaling, Danvers, MA) with protease and phosphatase inhibitors (Boston BioProducts, Boston, MA). Total lipid was extracted using the Folch method [18] and final triglyceride concentrations were determined using a spectrophotometric assay as previously described [19].

### Glucose Tolerance Test (GTT)

Glucose tolerance was determined after an overnight fast of mice at 5 and 9 months of age with 5 per group. Mice were weighed and injected intraperitoneal with 20% glucose (2g/kg) and blood glucose levels, collected from tail, were measured over a 120-minute period using a glucometer (Contour NEXT EZ, Bayer, Whippany, Germany). The area under curve (AUC) for each curve was determined and represented as AUC glucose (mmol X 120 min).

### Insulin Tolerance Test (ITT)

Insulin Tolerance was determined after an overnight fast of mice at 5 and 9 months of age with 5 per group. Mice were weighed and injected intraperitoneally with 0.75 Units/kg body weight and blood glucose levels, collected from tail, were measured over a 120-minute period using a glucometer (Contour NEXT EZ, Bayer, Whippany, Germany). The area under curve (AUC) for each curve was determined and represented as AUC glucose (mmol X 120 min).

### IGF-1 Levels

IGF-1 levels in the serum and tissues (liver, gastrocnemius, and brain cortex) were determined using Quantikine mouse IGF-1 immunoassay from R&D systems (MN, USA). Serum samples were diluted in calibrator diluent provided in the kit at 500-fold dilution and assayed according to the manufacturer’s instructions. Values are represented as pg of IGF-1 per ml of serum. For the tissue levels, IGF-1 was first isolated from its binding components by acid extraction using sodium acetate buffers (pH 3.6) before proceeding with the ELISA as described by Adams et al. [20]. Values are represented as pg of IGF-1 per 50 mg of tissue.

### FGF21 Levels

FGF-21 levels in the serum and liver were determined using Quantikine mouse/rat FGF-21 ELISA kit from R&D systems (MN, USA). Serum and liver homogenates were measured by a solid-phase ELISA technique according to the manufacturer’s instructions. Value are represented as pg/mg of tissue or pg/ml of serum of FGF-21.

### Adiponectin levels

Adiponectin levels in the gonadal white adipose tissue was determined using Quantikine mouse adiponectin/Acrp 30 ELISA kit from R&D systems (MN, USA). Liver homogenates were measured by a solid-phase ELISA technique according to the manufacturer’s instructions. Value are represented as pg/mg of tissue of Adiponectin.

### Real-time PCR

The levels of specific mRNA transcripts of genes involved in hunger/satiety, inflammation and fatty acid metabolism were measured by real-time PCR in the hypothalamus and gonadal white adipose tissues of litter expansion mice (n of 4-5/group). Briefly, RNA was isolated using the RNeasy kit from Qiagen (Germantown MD, USA). The first strand cDNA was synthesized from 1µg RNA using random primers (Promega, Madison, WI, USA) and purified using the QIAquick PCR purification kit (Qiagen, Germantown, MD, USA). Expression of some of the candidate genes (IL-6, TNF-α, MCP-1, FAS, ACC, CPT-1, MCAD, LCAD, β-Actin) were quantified using real-time PCR with SYBR green and the primer sequences are given in Table 1. Expression of Pomc (Mm00435874_m1), Npy (Mm01410146_m1), AgRP (Mm00475829_g1), CRH (Mm01293920_s1) and β-Actin (Mm02619580_g1) in the hypothalamus was done using specific Taqman probe sets. The gene transcripts were normalized to β-actin. Relative gene expression was quantified as comparative ct analysis using the 2^-ΔΔct^ analysis method with β-actin as endogenous control. One-way ANOVA design with Tukey’s multiple test correction was used to statistically analyze individual samples.

**Table 1:**
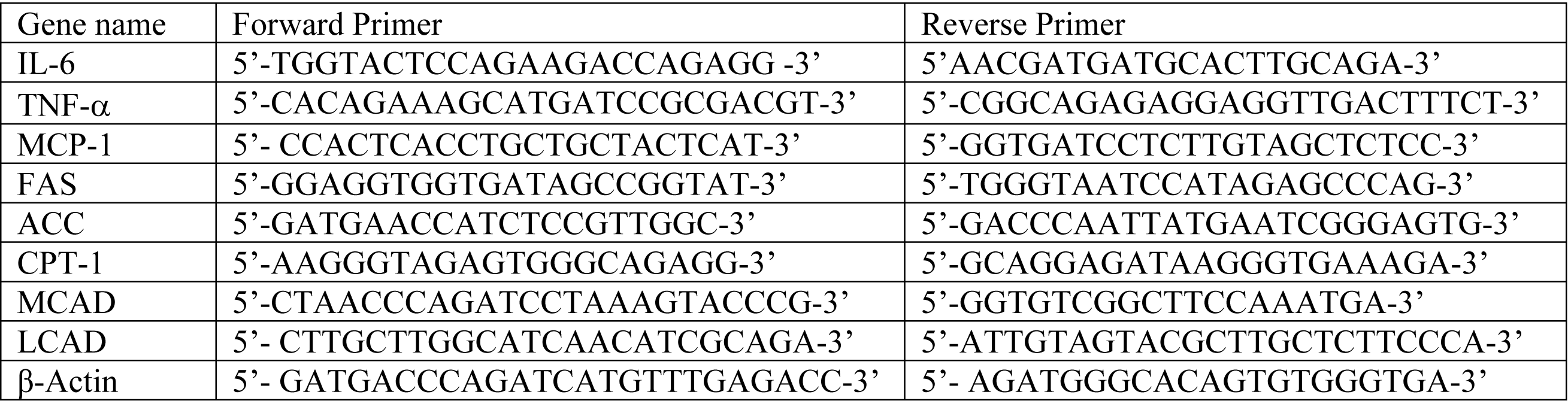
Primer Sequence.

## Results

### Effect of litter expansion on body weight and body composition

The average litter size for C57BL/6J mice is 6; therefore, we studied the effect of litter size expansion from 6 pups/litter (LS6) to litter sizes of 8 (LS8), 10 (LS10) and 12 (LS12), i.e., we expanded the litter size by 33%, 66% and 100%, respectively. We also reduced the litter size by ∼30% to 4 pups/litter (LS4) to study the effect of reduced litter size on various parameters. We first determined the effect of litter expansion on the development of sexual maturity in the female mice as measured by age of vaginal patency. We found that all female mice studied from litter sizes ranging from 4 to 10 showed vaginal patency at a similar age, e.g., ∼32 days of age. However, only 44% of the female mice from 12 pups/litter showed vaginal opening by 32 days of age. This suggests a delay in sexual maturity in LS12 female mice which has been correlated to an increase in lifespan [17].

Because Sun et al. [15] showed that pre-weaning food restriction reduced the size and body weight of the mice at weaning and maintained reduced body weight throughout their lifespan, we followed the changes in body weight and composition through 15 months of age. Fig 1 shows the body weight of both male and female crowded litter animals at 1, 5, 9, 11 and 15 months of age. LS12 male and female mice showed a significantly reduced body weight at one month of age compared to control, LS6 mice, and they maintained reduced body weight through 15 months of age. In males, LS8, LS10, and LS12 mice showed significant body weight reduction at 15 months of age (Fig1). The effect of litter expansion on body weight was more pronounced in male mice especially at later ages such that at 15 months of age, e.g., the LS12 males showed ∼14% decline in body weight compared to LS6 males while females LS12 did not show a significant difference in body weight compared to LS6 females (Fig1). It should be noted that we observed no significant difference in the body weight of LS4 and LS6 mice at any age or sex.

**Fig 1.**
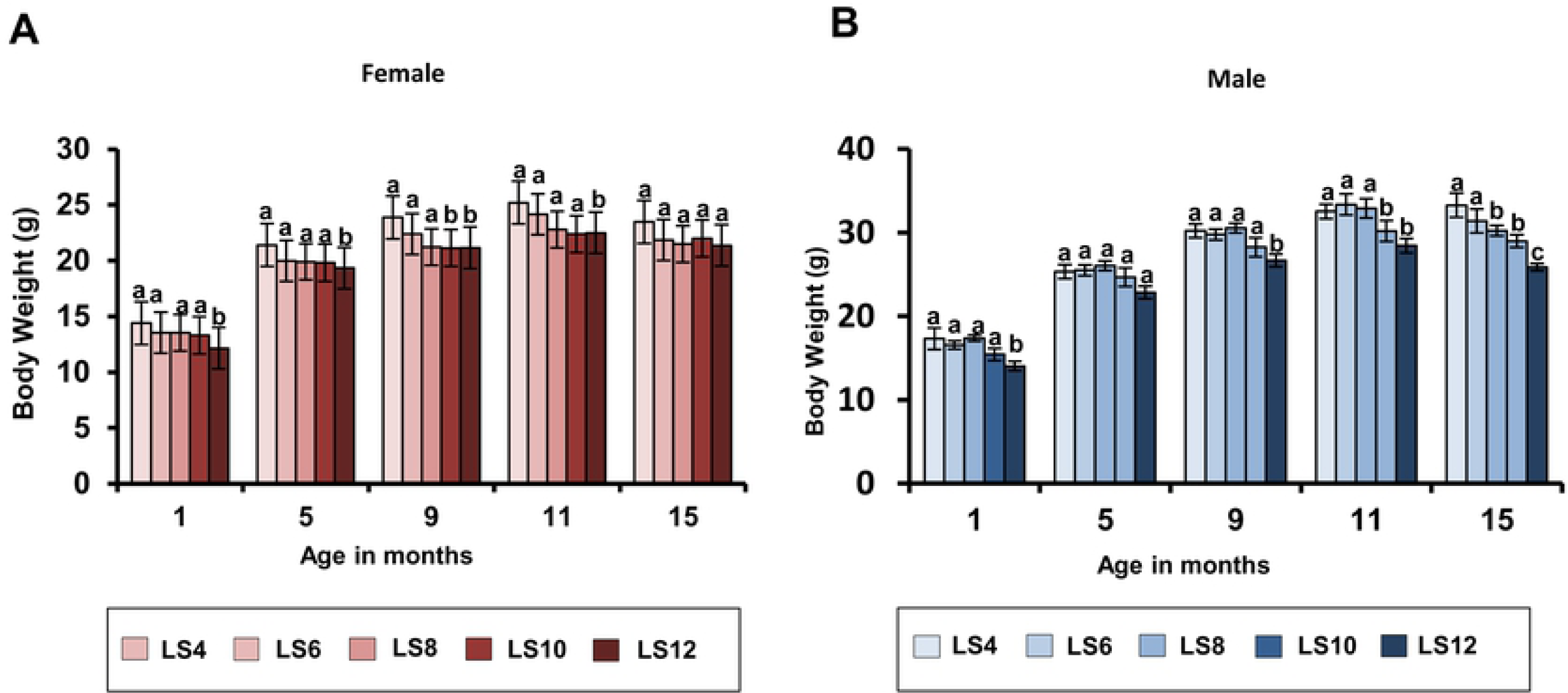
Effect of litter expansion on body weight. The body weight of female and male mice from different litter sizes (LS4, LS6, LS8, LS10, LS12) were measured at 1, 5, 9, 11, and 15 months of age. Data represented are the mean ± SEM from 5-12 mice per group and were statistically analyzed by one-way ANOVA with Tukey’s multiple correction test. Different alphabetic letters show significant difference (P < 0.05) for each sex whereas, same alphabets denotes no significance.

Body composition of the crowded litter mice was measured at 15 months of age. Fig 2A and 2B show the lean body mass (LBM) and fat mass measured by NMR along with the weights of liver and gastrocnemius muscle. Both female and male mice showed statistically significant difference between groups only in their total fat mass and showed no difference in their LBM and weights of liver and gastrocnemius. The male and female litter expanded mice showed a major sex difference in their level of reduction in total fat content. Female LS8, LS10, and LS12 mice showed ∼17% reduced total fat mass compared to LS6 mice, whereas the male LS10 and LS12 mice showed a much more robust (50 and 60% respectively) reduction. Again, we observed no significant difference in body composition of the L4 and LS6 mice. From the body composition data, it is evident that the reduced body weight observed in the crowded litter mice at 15 months of age is due primarily to a reduction in total fat mass and this decrease was more pronounced in male mice.

**Fig 2.**
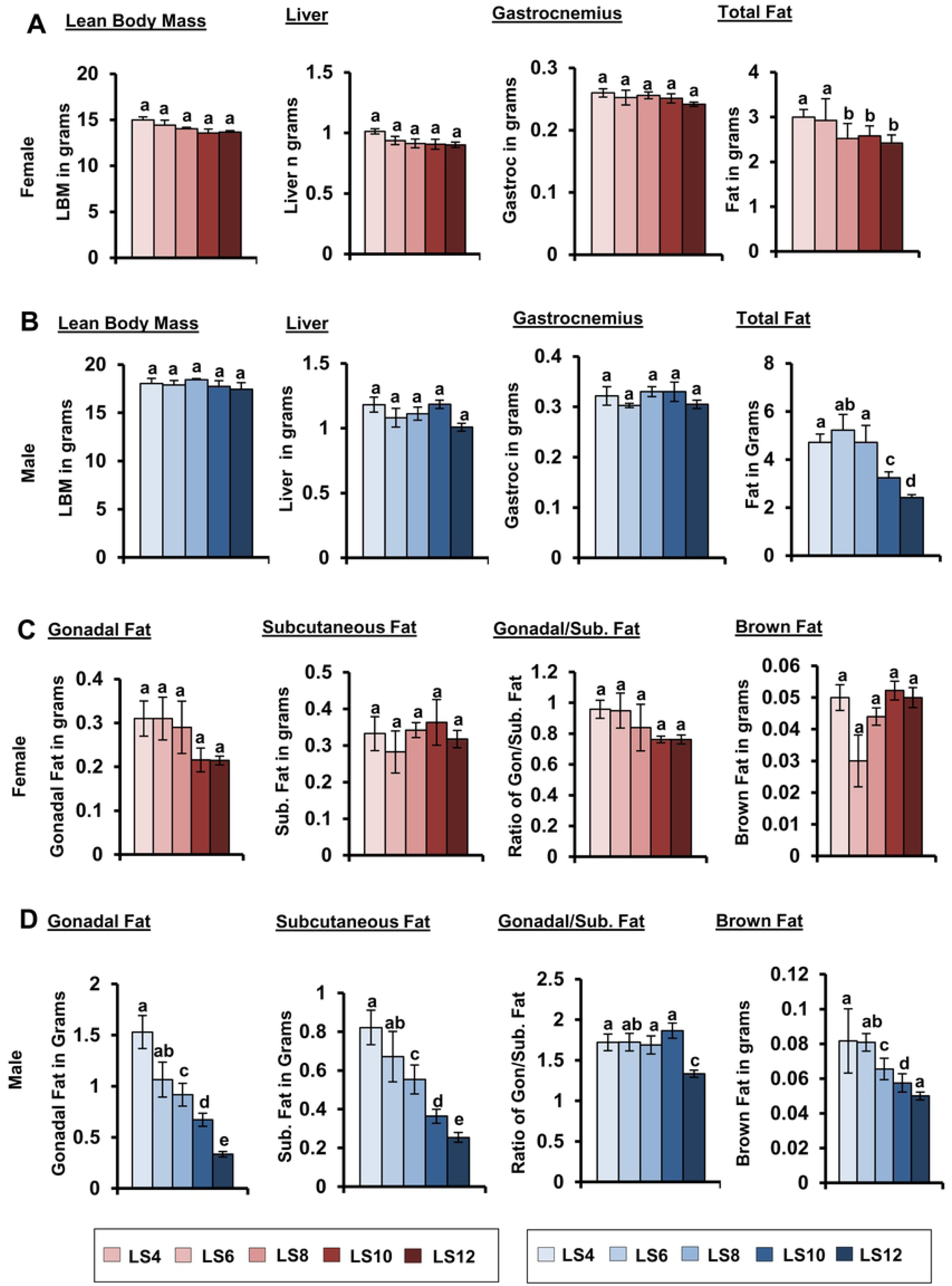
Effect of litter expansion on body composition and fat depots in 15-month-old mice. The lean body mass, total fat, mass of liver, and gastrocnemius in (A) female and (B) male mice and mass of different fat depots: gonadal, subcutaneous and brown fat in (C) female and (D) male mice. Data represented are the mean ± SEM from 5 mice per group and were statistically analyzed by one-way ANOVA with Tukey’s multiple correction test. Different alphabetic letters show significant difference (P < 0.05) for each sex whereas, same alphabets denotes no significance.

Because the different fat depots have different phenotypic effects, we measured the weights of several fat depots in the mice at 15 months of age: gonadal fat, posterior subcutaneous fat (which includes dorsolumbar, inguinal and gluteal fat), and brown fat (Fig 2 C and 2D). Visceral fat, which includes gonadal fat, is associated with metabolic dysfunction and insulin resistance, whereas subcutaneous fat is considered to be protective against the development of insulin resistance [21]. Litter expansion did not have any significance effect on the fat depots of female mice, although the gonadal fat showed a trend in reduction in the LS10 and LS12 mice (Fig 2C). On the other hand as shown in Fig 2D, LS10 and LS12 mice showed a significant decline in all three fat depots with the LS12 mice showing a greater decline than the LS10 mice compared to LS6 male mice, e.g., a 75% vs 25% decrease in gonadal fat; a 64% vs 50% decrease in subcutaneous fat, and a 38% vs 25% decrease in brown fat. We also measured the ratio of gonadal fat to subcutaneous fat (Fig 2C and 2D) because the adipose tissue in the subcutaneous area differs from the visceral fat in cell size, metabolic activity potential role in insulin resistance [22]. The visceral fat is considered to be a “bad fat” and more pathogenic towards obesity induced insulin resistance whereas the subcutaneous fat is the “good fat” and is considered beneficial [22]. Male LS12 mice showed a significantly lower ratio of gonadal to subcutaneous fat (∼29%), indicating that they have proportionally more subcutaneous fat (good fat) than gonadal fat (bad fat) (Fig 2D). Female LS10 and LS12 mice also show a trend to a lower (∼17%) ratio of gonadal to subcutaneous fat; however, this decrease was not statistically significant (Fig 2C). It should be noted that we observed no difference in the fat depots for LS4 and LS6 male or female mice. From these data, it is evident that litter expansion exhibits a sexual divergence in adiposity at 15 months of age in response to a manipulation that occurred just during the prenatal period. Because litter expansion induced significant changes in the fat depots in male mice, we also measured the triglyceride content of liver in these mice because fatty liver can lead to negative outcomes. There was no difference in the triglyceride content of LS12 and LS6 male or female mice as shown in S1 Fig.

Because the neural circuits (POMC, NPY and AgRP) in the arcuate nucleus (ARC) of the hypothalamus are involved in appetite regulation (e.g., POMC is anorexigenic and NPY and AgRP is orexigenic) and develop primarily during the first 3 weeks of postnatal life in rodents [23], we were interested determining if the changes in adiposity induced by litter expansion arose from alterations in these neural circuits. Therefore, we measured the transcript levels of Pomc, AgRP, NPY and CRH in the hypothalamic of 15-month-old male and female mice. As shown in the S2 Fig, we did not observe any significant difference in the levels of the transcripts for any of the hypothalamic genes (except pomc in male LS10) suggesting there were no changes in the markers for appetite control and stress at least in the transcript level.

### Effect of litter expansion on fat metabolism, cytokines and adipokines

To gain an insight into the effect of litter expansion on the fatty acid metabolism, we measured the expression of genes involved in fatty acid synthesis and oxidation. Dietary restriction has been shown to increase the mRNA levels of key genes involved in fatty acid synthesis [e.g., Fatty Acid Synthase (FAS) and Acetyl CoA Carboxylase (ACC)] and fatty acid oxidation [e.g., Medium Chain Acyl-CoA dehydrogenase (MCAD) and Long Chain Acyl-CoA dehydrogenase (LCAD)] in white adipose tissue [24]. Changes in these key genes regulates fatty acid metabolism such that increases in FAS and ACC increase fatty acid synthesis and increases in MACD and LCAD increase β-oxidation of middle and long chain fatty acids. Fig 3A shows the mRNA levels of all the four genes in the gonadal tissue of the male and female mice. Litter expansion in the female mice showed a trend for reduced expression of these genes; however, the differences were not statistically significant because of the large animal to animal variation. In contrast, male LS10 and LS12 mice showed a 100% increase in LCAD and a significant decrease (75%) in FAS gene expression (Fig 3B) indicating a potential increase in fatty acid oxidation and reduction in fatty acid synthesis which in turn could explain the decrease in gonadal adipose tissue weight (Fig 2D).

**Fig 3.**
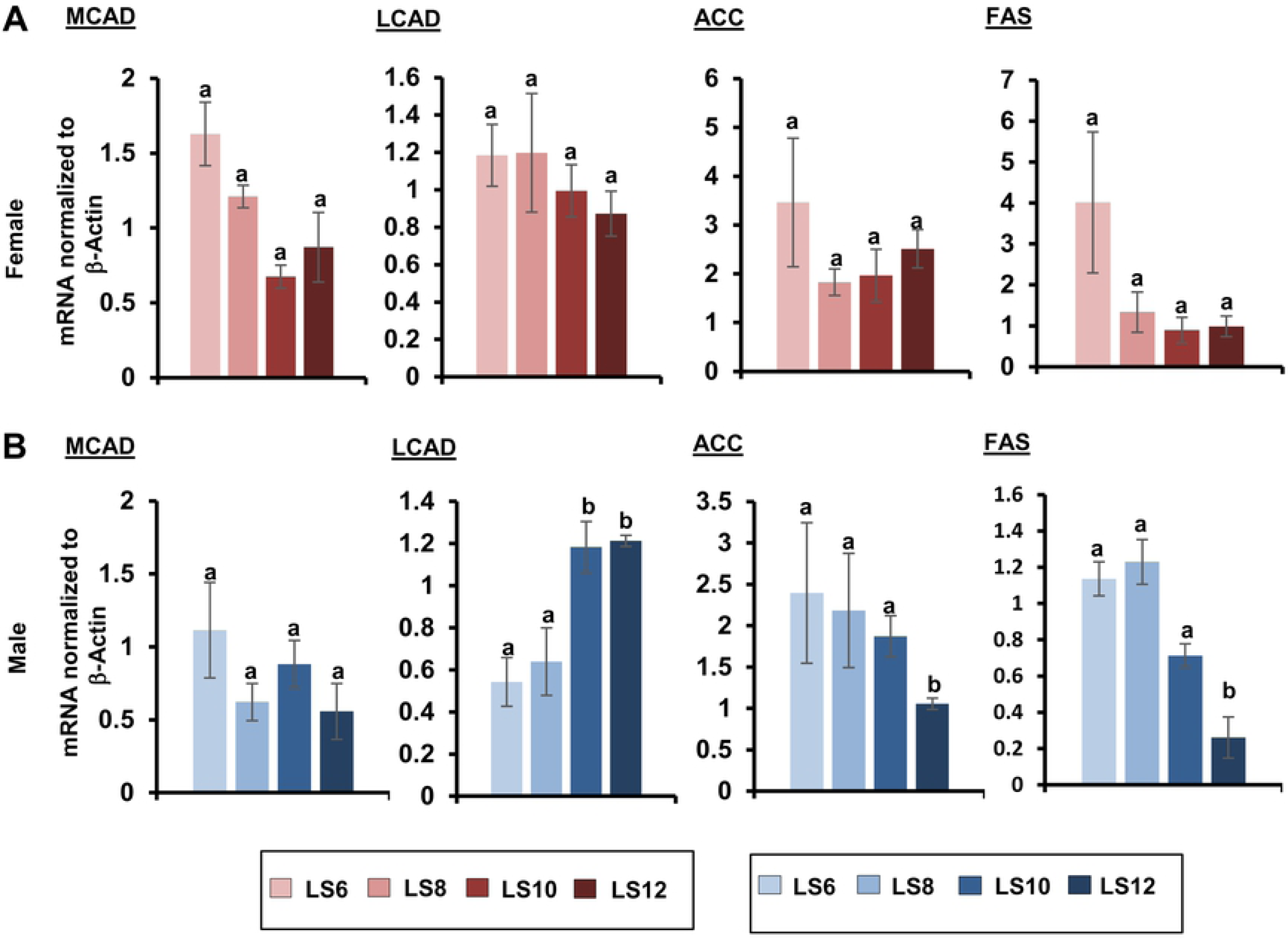
Effect of litter expansion on expression of genes involved in fatty acid metabolism. The mRNA levels of Medium Chain Acyl-CoA dehydrogenase (MCAD) and Long Chain Acyl-CoA dehydrogenase (LCAD), genes involved in fatty acid break down, and Acetyl CoA Carboxylase (ACC) and Fatty Acid Synthase (FAS) genes involved in biosynthesis were measured in the gonadal fat of female (A) and male (B) mice from various litter sizes at 15 months of age. Data represented are the mean ± SEM from 3-5 mice per group and were statistically analyzed by one-way ANOVA with Tukey’s multiple correction test. Different alphabetic letters show significant difference (P < 0.05) for each sex whereas, same alphabets denotes no significance.

White adipose tissue is a major endocrine and secretory organ capable of releasing a variety of pro-inflammatory factors (e.g., IL-6, TNF-α and MCP-1) and changes in fat mass has been shown to affect the production pro-inflammatory cytokines [24]. Therefore, we measured the effect of litter expansion on the expression of IL-6, TNF-α and MCP-1 in gonadal fat from the male and female mice. As shown in Fig 4A, litter expansion had no significant effect on the expression of IL-6, TNF-α or MCP-1 in female mice. However, in male mice, litter expansion had a significant effect in the LS12 male mice. Compared to the other groups of mice, LS12 mice showed a significant increase (∼2.5-fold) in the levels of IL-6 and MCP-1 mRNA (Fig 4B), and the transcript levels of TNF-α tended to be higher (2-fold) than the control, LS 6 male mice; however, this increase was not statistically significant.

**Fig 4.**
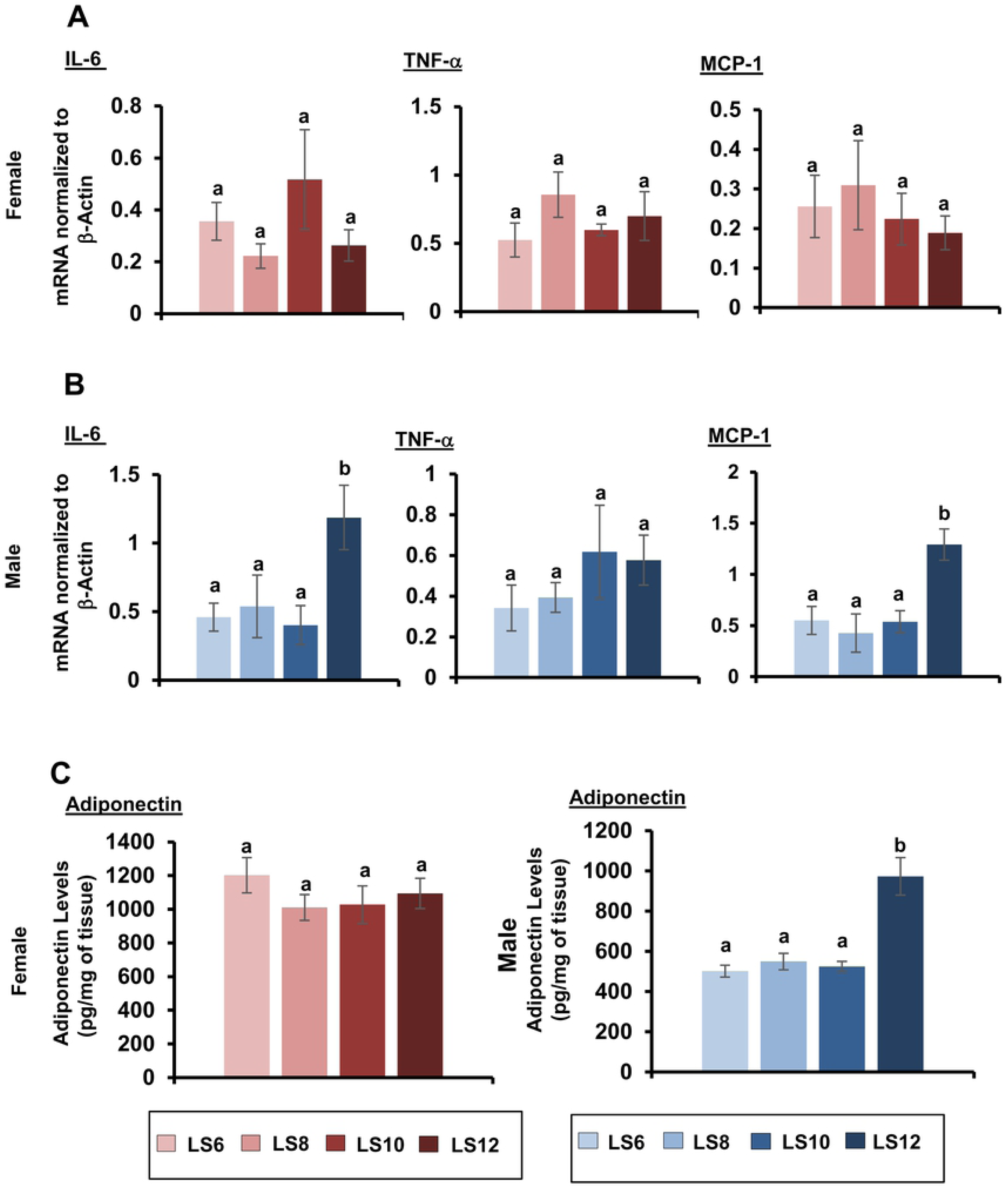
Effect of litter expansion on expression of genes involved in inflammation and adiponectin levels. Levels of mRNA of genes involved in inflammation (IL-6, TNF-α, MCP-1) were measured in the gonadal fat of (A) female and (B) male mice from various litter sizes (LS6, LS8, LS10 and LS12 pups/litter) at 15 months of age. Protein levels of adiponectin (C) in the gonadal fat of female and male mice from various litter sizes (LS6, LS8, LS10 and LS12 pups/litter) were measured at 15 months of age. Data represented are the mean ± SEM from 3-5 mice per group and were statistically analyzed by one-way ANOVA with Tukey’s multiple correction test. Different alphabetic letters show significant difference (P < 0.05) for each sex whereas, same alphabets denotes no significance.

White adipose tissue also secretes adiponectin which is important in adipocyte differentiation and can also function as an anti-inflammatory factor [25, 26], and dietary restriction has been shown to increase circulating adiponectin levels and increase adiponectin transcript levels in white adipose tissue [24, 27]. We measured the effect of litter expansion on levels of adiponectin protein in the gonadal tissue of the male and female mice. As shown in Fig 4, female mice show no difference in the levels of adiponectin in gonadal fat in any of the groups. In male mice, the LS12 group showed a significant (94%) increase in adiponectin levels compared to the other three groups (Fig 4C). Interestingly, the levels of adiponectin in tissues of the LS6, LS8, and LS10 male mice were significantly (50%) lower than the female mice except for male LS12 mice, which was in a similar range as observed in the females mice (Fig 4C). Our data are comparable to other studies showing that female mice have increased adiponectin levels compared to male mice [28].

### Effect of litter expansion on insulin sensitivity

Changes in adiposity have been shown to effect glucose metabolism and insulin sensitivity, e.g., visceral adipose tissue accumulation is associated with the development of insulin resistance [29, 30]. Aging is also associated with an increase in bodyweight, fat mass and insulin resistance [31], and anti-aging interventions such as dietary restriction and dwarfism increase insulin sensitivity in mice [32, 33]. Therefore, we were interested in studying the effect of litter expansion on insulin sensitivity by measuring glucose and insulin tolerance at 5 and 9 months of age in male and female mice (Fig 5). No difference in fasting blood glucose levels were observed between any of the groups of mice at either age (data not shown). Litter expansion had no effect of glucose tolerance in both male and female mice at 5 months of age. At 9 months of age, glucose tolerance was improved (∼18%) for the LS10 and LS12 mice compared to the other three groups for both male and female mice, i.e., litter expansion had a similar effect of glucose tolerance in male and female mice. However, we observed a major sex difference in insulin tolerance as shown in Fig 5. In female mice, litter expansion had no effect on insulin tolerance. In contrast, insulin tolerance in male mice was significantly improved at both 5 and 9 months of age in the LS10 and LS12 mice compared to the other three groups. For example, insulin tolerance was improved ∼9% at 5 months of age and ∼22% at 9 months of age. It should be noted that we observed no difference in either glucose or insulin tolerance in either male or female mice for the LS4 and LS6 mice.

**Fig 5.**
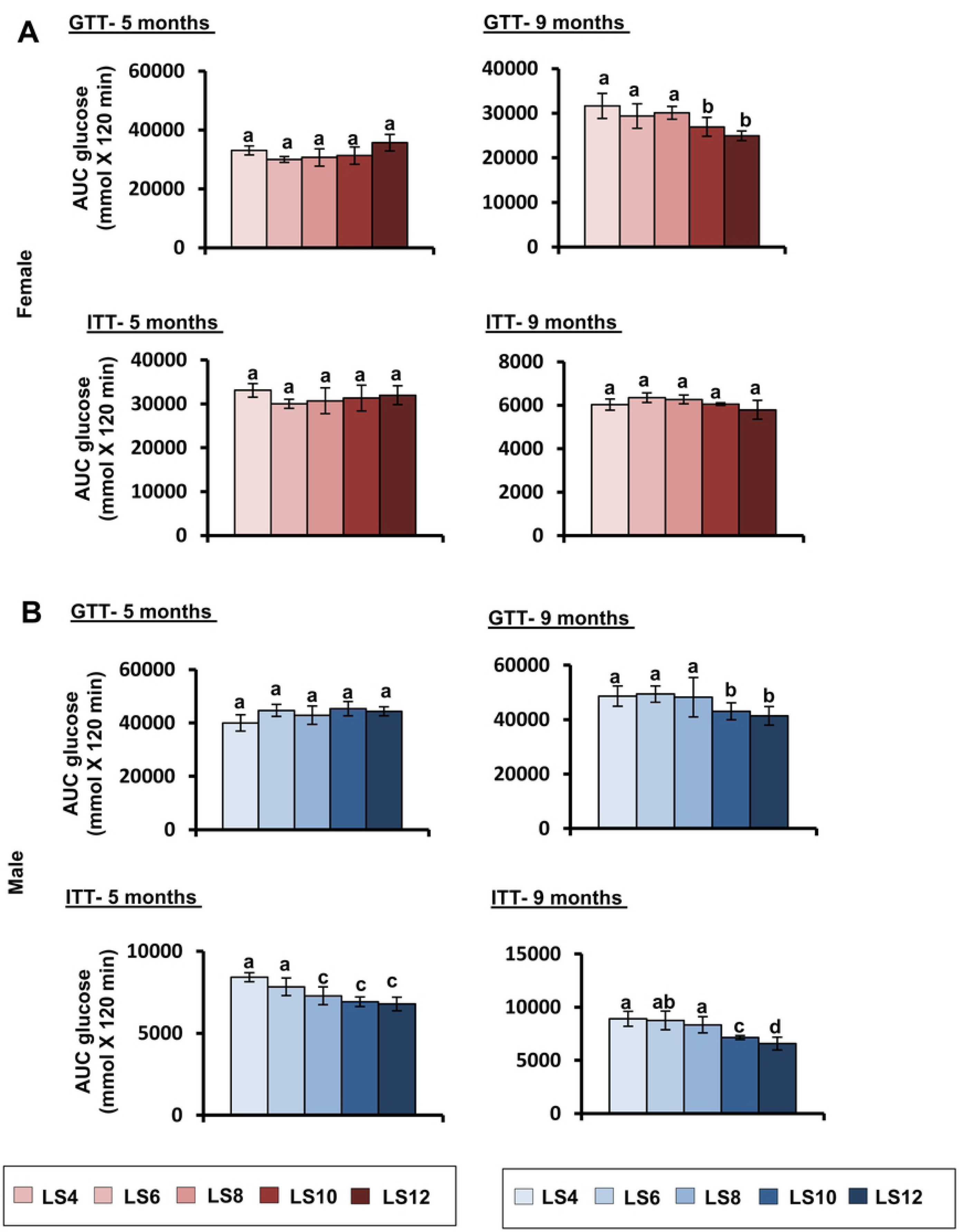
Effect of litter expansion on glucose homeostasis. Glucose tolerance (GTT) and insulin tolerance (ITT) were measured in female (A) and male (B) mice as described in the methods. The data are expressed as the area under the curve (AUC) for mice at 5 and 9 months of age. Data represented are the mean ± SEM from 5 mice per group and statistically analyzed by one-way ANOVA with Tukey’s multiple correction test. Different alphabetic letters show significant difference (P < 0.05) for each sex whereas, same alphabets denotes no significance.

### Effect of litter expansion on IGF-1 and FGF-21 expression

IGF-1 a neurotrophic factor that is primarily produced in the liver and secreted into the plasma. Reduced circulating IGF-1 levels have been shown to be associated with increased lifespan in dwarf mice [34] and dietary restricted mice [20, 35]. We first measured the circulating levels of IGF-1 in the serum of 15-month-old male and female mice. As shown in Fig 6A, IGF-1 levels were significantly reduced (∼25-30%) in the serum of both female and male LS8, LS10, and LS12 mice compared to LS6, control mice. As would be predicted, the decrease in circulating IGF-1 was directly correlated with reduced levels of IGF-1 protein in the liver of the LS8, LS10, and LS12 mice (Fig 6B). Because IGF-1 is produced by other tissues, we also measured the levels of IGF-1 protein in the gastrocnemius and brain cortex of 15-months old mice. The levels of IGF-1 protein in the gastrocnemius was reduced in the LS8, LS10, and LS12 mice compared to LS6 mice for both male and female mice (Fig 6C). However, the effect of litter expansion on IGF-1 protein levels in the cortex was sexually divergent. IGF-1 levels were the same in all four groups of female mice; however, IGF-1 levels in the brain cortex of male LS8, LS10, and LS12 mice were significantly lower (∼75-95%) than the LS6 mice (Fig 6D). To determine if litter expansion reduced IGF-1expression at the level of transcription, we measured the levels of IGF-1 mRNA in liver, gastrocnemius, and cortex of all four groups of male and female mice. As shown in S3 Fig, we observed no significant change in IGF-1 mRNA levels in any of the groups of either male or female mice. In other words, the reduction in IGF-1 levels was not due to reduced transcription.

**Fig 6.**
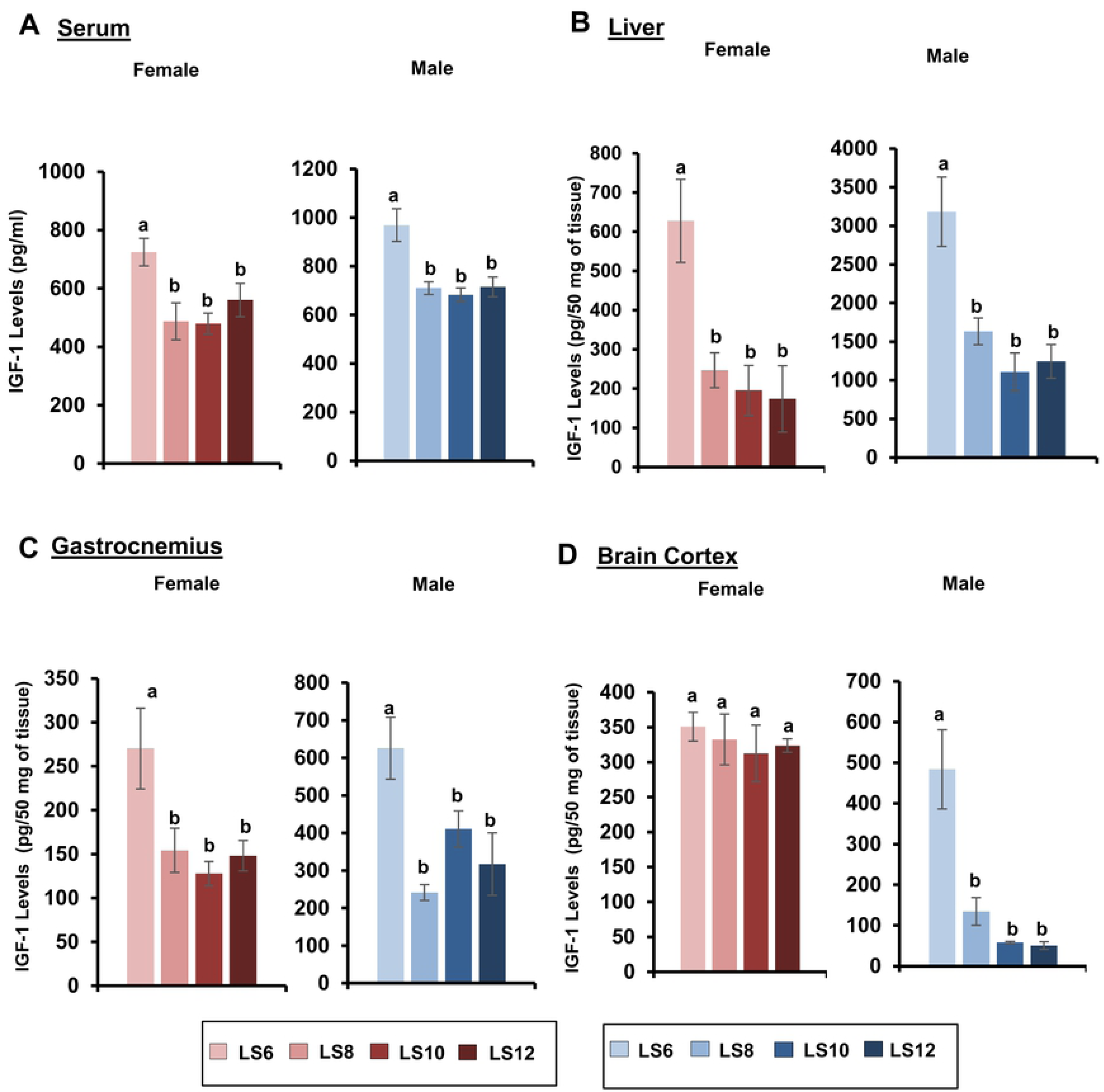
Effect of litter expansion on IGF-1 levels. The protein levels of IGF-1 protein were measured in serum (A), liver (B), gastrocnemius (C), and cortex (D) of 15-month-old male and female mice as described in the Methods. Data represented are the mean ± SEM from 3-5 mice per group and were statistically analyzed by one-way ANOVA with Tukey’s multiple correction test. Different alphabetic letters show significant difference (P < 0.05) for each sex whereas, same alphabets denotes no significance.

Fibroblast growth factor-21 (FGF21) is a metabolic hormone produced predominantly in the liver and secreted into the circulation similar to IGF-1. FGF-21 has been shown to have a pronounced effect on glucose and lipid metabolism. We first measured the effect of litter expansion on the levels of FGF21 in liver of crowded litter of the male and female mice. Female mice did not show any change in liver FGF21 levels (Fig 7A). In contrast, male LS8, LS10, and LS12 mice showed a 100% Increase in the level of FGF21 compared to LS6 mice (Fig 7A). We then determined if the changes in FGF21expression in liver of male mice resulted in changes in circulating FGF21. As shown in Fig 7B, serum FGF21 levels were also significantly increased (100-150%) in the LS8, LS10, and LS12 mice compared to the LS6 mice.

**Fig 7.**
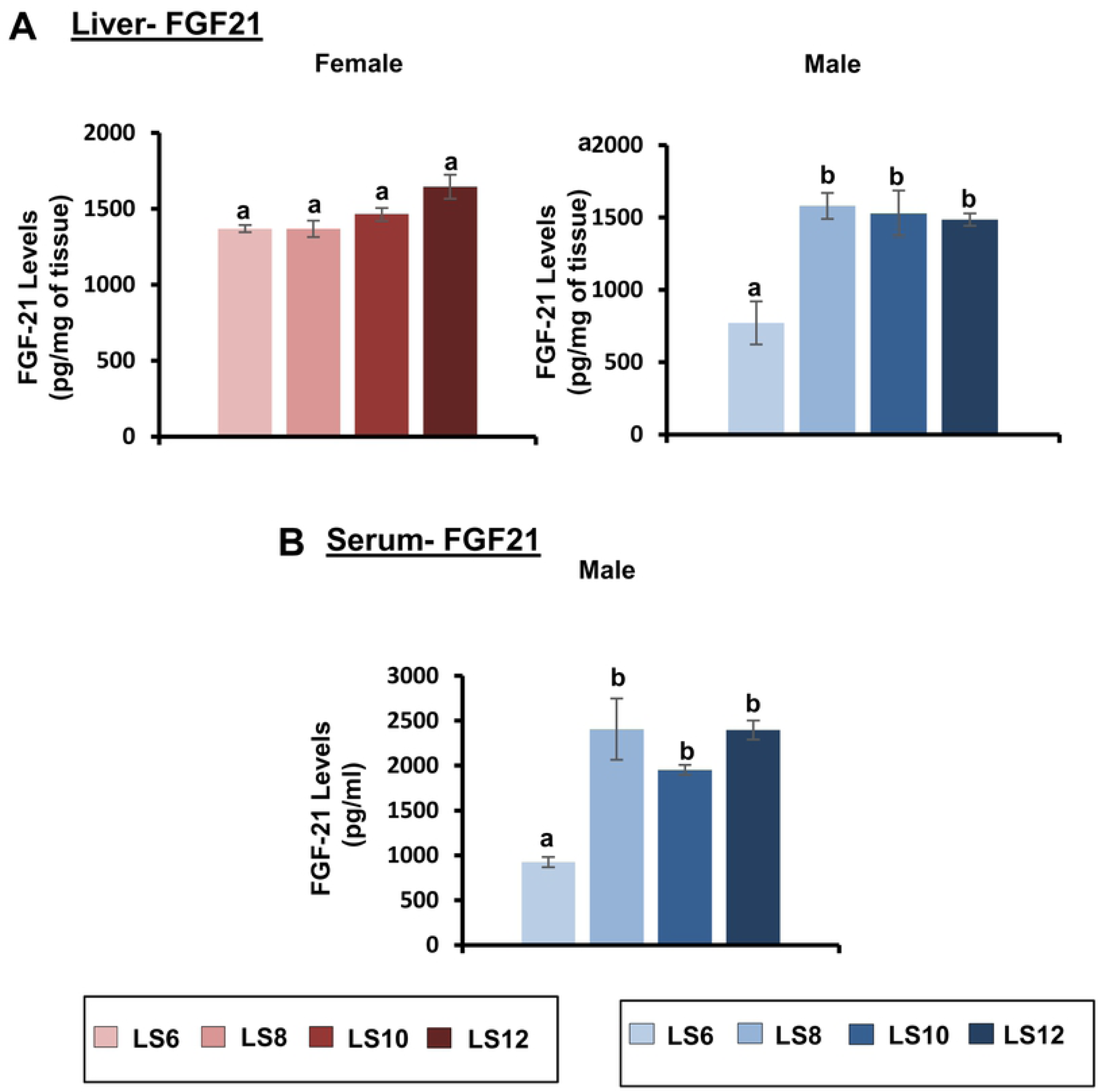
Effect of litter expansion on FGF21 levels. The protein levels of FGF21 in the liver (A) of female and male mice and serum (B) of male mice were measured at 15 months of age. Data represented are the mean ± SEM from 3-5 mice per group and were statistically analyzed by one-way ANOVA with Tukey’s multiple correction test. Different alphabetic letters show significant difference (P < 0.05) for each sex whereas, same alphabets denotes no significance.

## Discussion

The classical study by McCay et al. [36] reported that a dramatic reduction in food consumption initiated at weaning in rats resulted in a 50% increase in lifespan. Multiple studies over the following 5 decades clearly demonstrated that reducing food intake by 30 to 40% (dietary restriction) increased the lifespan of rats and mice by 20 to 30%, in addition to improving health and reducing pathological lesions [37]. McCay et al. [36] initially proposed that preventing growth was key to the effect of restricting the food intake on longevity. This view was held until the 1980s when it was shown that dietary restriction started after growth, e.g., at 6-months of age in rats [38] or 12-months of age in mice [39] increased lifespan significantly. These findings led investigators to primarily study the effect of dietary restriction after mice or rats had reached sexual maturity, e.g., 2 to 6 months of age. However, Yu et al. [38] showed that dietary restriction initiated in rats shortly after weaning for only 18 weeks resulted in a significant increase in lifespan, suggesting that dietary restriction for a limited time early in life could have a long-term effect on the rate of aging. This concept was reinforced when Sun et al. [15] showed that food restriction during the lactation period implemented by litter expansion increased the mean and maximal lifespan of mice.

The purpose of this study was to comprehensively characterize the long-term effects of litter expansion in mice. Our study differed in several ways from the previous reports with UM-HET3 mice where the litters of the mice were culled to 8 pups/litter and then either 4 or 7 additional pups were added to the litters giving litter sizes 12, and 15 compared to the control of 8 pups/litter [15, 16]. It should be noted that the average litter size of UM-HET3 mice is ∼10 pups. We used C57BL/6J mice, which have an average litter size of ∼6 pups, and generated litter sizes of 6, 8, 10 and 12, i.e., we studied the effect of increasing the litter size from 33 to 100% over the normal litter size of 6 pups. We also studied the effect of reducing the litter size to 4 pups per litter as a potential model of over-nutrition. We found no short or long-term effects of reducing the litter size from 6 to 4 mice; however, we did observe significant short- and long-term effects of litter expansion, which were found to elicit sexually divergent responses.

As previously reported [15], we observed a significant decrease in the body weight of the LS12 mice that was detected at one month of age and was maintained until 15 months of age in both male and female mice. However, the effect of litter expansion on body weight was much greater in male mice such that the reduction in body weight became more pronounced with age. At 15 months of age, LS8, LS10, and LS12 male, all mice showed a significant reduction in body weight compared to the control (LS6) with the LS12 male mice showing a ∼20% reduction in body weight. Sadagurski et al. [16] reported that the decrease in body weight at 2 months of age due to litter expansion in UM-HET3 mice arose from a decrease in body fat with no change in lean body weight in both male and female UM-HET3 mice. We also found that the reduction in body weight with litter expansion was accompanied by a reduction in body fat in both males and females at 15 months of age with no change in lean body weight. However, the decrease in total fat was much greater in male mice, e.g., a 50% decrease in LS12 males compared to a modest 15% in LS12 females. Our study is the first to determine how litter expansion affects specific fat depots (gonadal, subcutaneous and brown fat). We found that sex differences in fat depot masses become more pronounced at 15 months of age. In female mice, a decreasing trend was only observed in gonadal fat, but was not statistically significant. In contrast, a significant reduction in gonadal, subcutaneous and brown fat was observed in male LS10 and LS12 mice compared to LS6 mice. Gonadal fat showed the greatest decrease (over 85% for LS12 vs LS6) and brown fat showed the smallest decrease (40% for LS12 vs LS6). Based on our data from the transcript levels of genes involved in fatty acid metabolism, it appears that litter expansion resulted in an increase in fatty acid oxidation gene, Long Chain Acyl-CoA dehydrogenase (LCAD) and a reduction in Fatty Acid Synthase (FAS) and Acetyl CoA Carboxylase (ACC) in male mice. Dietary restriction implemented in adult mice also shows a reduction in adiposity; however, the effect is quite different than we have observed with litter expansion. For example, 4-months of dietary restriction (40%) reduced the fat content in both male and female C57BL/6J mice equally (unpublished data), whereas reductions in adiposity by litter expansion is more robust in males.

The reduction in adiposity by dietary restriction has been correlated to reduced expression of pro-inflammatory cytokines and increased expression of adiponectin, which has anti-inflammatory effects [25, 26]. Litter expansion had no effect on the transcript levels of IL-6, TNF-α, or MCP-1 or the levels of adiponectin protein in gonadal fat of female mice. In contrast, the LS12 male mice showed a significant increase in the transcript levels of IL-6 and MCP-1 and adiponectin protein levels in gonadal fat. We were surprised to observe an increase in IL-6 and MCP-1 because they are considered to have detrimental effects and are correlated with reduced glucose and insulin tolerance, both of which were increased in the LS12 male mice. However, there are data suggesting that elevation in these markers can have beneficial metabolic effects [40, 41]. For example, increased levels of IL-6 has been shown to be correlated to improved glucose homeostasis [40, 42].

One of the hallmark characteristics of dietary restriction is improved glucose homeostasis, e.g., improved glucose tolerance and insulin sensitivity. Matyi et al. [24], showed that improved glucose tolerance could be observed in male C57BL/6J mice within 10 days of reducing (40% restriction) food consumption of 4-month-old mice and the improved glucose tolerance was maintained when DR was discontinued. In a later study we have observed no sex differences in either glucose tolerance or insulin sensitivity when dietary restriction was implemented in 6 weeks-old C57BL/6J mice (unpublished data). Sadagurski et al. [16] reported that litter expansion improved glucose tolerance and insulin sensitivity at 6 months of age in UM-HET3 male mice but not female mice. However, at 22 months of age, litter expansion improved glucose tolerance (insulin sensitivity was not measured) in the female mice. We observed a similar effect in C57BL/6J mice. At 5 months of age, females show no significant difference in either the glucose or insulin tolerance. At 9 months of age, the LS10 and LS12 female mice showed a significant improvement in glucose tolerance compared to LS6 mice. In male mice, we also observed that litter expansion had no significant difference on glucose tolerance at 5 months of age. However, the LS10 and LS12 male mice showed significant increase in insulin tolerance compared to the LS6 male mice. At 9 months of age, LS10 and LS12 male mice showed significant improvement in both glucose tolerance and insulin tolerance. Collectively, these observations indicate that litter expansion promotes a longer window of metabolic plasticity in both sexes with advancing age.

We were interested in studying the long-term effects of litter expansion on IGF-1 because changes in IGF-1 expression have been associated with longevity. For example, early studies showed that dietary restriction reduced circulating IGF-1 levels in rats [43] and this has been observed in mice [44]. Dwarf mice, which have increased lifespan, also have lower levels of IGF-1. In addition, mice heterozygous for IGF-1 receptor have been reported to have a ∼30% increase in lifespan [45]; however, more recent studies show only a modest increase in lifespan of ∼6% in *Igf1r*^+/-^ mice [46, 47]. Furthermore, Ashpole et al. [48], showed that IGF-1 levels during the developmental phase plays an important role in late-life healthspan and lifespan, and Sun et al. [15] reported that litter expansion reduced serum IGF-1 levels in weanling mice but did not observe a sex effect on the levels at this age. We found that serum levels of IGF-1 were significantly reduced (∼25-30%) by litter expansion in both male and female mice at 15 months of age. Interestingly the reduction in serum IGF-1 levels was similar in LS8, LS10, and LS12 mice, i.e., the reduction in IGF-1 levels did not increase when the litter size was increased over 8 pups/litter. As would be expected, reduction in circulating IGF-1 levels was correlated to a dramatic reduction (over 50 %) in IGF-1 protein levels in the liver. We also studied the effect of litter expansion on the endogenous expression of IGF-1 in skeletal muscle and brain. A decrease in IGF-1 protein levels were observed in the gastrocnemius from both male and female LS8, LS10, and LS12 mice. However, when we measured the levels of IGF-1 protein in brain cortex we found a dramatic sex effect. Litter expansion had no effect on IGF-1 protein levels in the cortex of female mice. However, male mice showed a very dramatic decrease (>80%) in LS8, LS10, and LS12 male mice compared to LS6 male mice. Interestingly, litter expansion had no effect on IGF-1 mRNA levels in any of the tissues studied; therefore, it appears that the effect of litter expansion on IGF-1 occurs post-transcriptionally. Interestingly, IGF-1 levels have been shown to be influenced by various microRNA [49, 50], e.g., miR-1 and miR-206 have been shown to target the 3-untranslated region of the IGF-1 mRNA molecule and reduce its expression [49]. Hence, more work is warranted to determine if the levels of IGF-1 are regulated by the microRNA’s in the litter expansion model.

Fibroblast growth factor-21 (FGF21) is considered to be a starvation hormone that stimulates gluconeogenesis, fatty acid oxidation, and ketogenesis as an adaptive response to fasting and starvation [51]. Overexpression of FGF21 has been shown to improve insulin sensitivity, blood glucose, lipid profile and body weight in obese and diabetic animal models [51] and increase the lifespan of normal mice [52]. Kuhla et al. [53] reported that dietary restriction increased plasma levels of FGF-21. In contrast, Miller et al. [54] reported that DR resulted in a decrease in plasma FGF-21 levels. We found that litter expansion resulted in an increase (∼100%) in the levels of FGF-21 protein in the livers of male mice but not female mice. The increased levels of FGF-21 in liver was associated with an increase in serum FGF-21 levels in male LS8, LS10, and LS12 mice. The litter expansion had no effect on the mRNA levels of FGF-21 in the livers of male or female mice. Thus, the increase in FGF-21 protein levels observed with litter expansion in male mice occur post-transcriptionally.

In summary, our study confirms data from the previous reports, showing that litter expansion can have long-term effects on body weight, adiposity, and glucose homeostasis in C57BL/6J mice in addition to other previously used strains of mice. However, we show for the first time that major sex differences occur in the long-term effects of litter expansion in mice with greater effects in males than females, e.g., adiposity, glucose homeostasis. In addition, we showed that litter expansion has a long-term effect on circulating levels of IGF1 and FGF-21, and there are major sex differences in brain levels of IGF1 and liver levels of FGF-21. Interestingly, dietary restriction, which is implemented in rodents between 2 to 6 months of age when mice are sexually mature, has similar effects in male and female C57BL/6J mice, e.g., the increase in lifespan is ∼20% in both male and female mice [55] and changes in growth are similar in males and females and we have further found that the improvement in glucose tolerance and insulin sensitivity is similar in male and female C57BL/6J mice (Unpublished data). Thus, the sexually divergent effect of dietary restriction implemented during the first weeks after birth by litter expansion is likely due to the dramatic changes in sexual maturation which occur in the first three weeks of life in mice [56] compared to when dietary restriction is implemented after the mice are sexually mature and are adults.

## Acknowledgements

None to report.

## Supporting information

**S1 Fig. Effect of litter expansion on liver triglyceride content**. The triglyceride content of the liver obtained from LS6 and LS12 were measured in female and male mice at 15 months of age. The liver triglyceride content data represented are the mean ± SEM from 5 mice per group and were statistically analyzed by two-sided t-test. Different alphabetic letters show significant difference (P < 0.05) for each sex whereas, same alphabets denotes insignificance.

**S2 Fig. Effect of litter expansion on expression of genes in hypothalamus**. Levels of mRNA of Pomc, Npy, AgRP and CRH genes were measured in the hypothalamus of female and male mice from various litter sizes (LS6, LS8, LS10 and LS12 pups/litter) at 15 months of age. Data represented are the mean ± SEM from 3-5 mice per group and were statistically analyzed by one-way ANOVA with Tukey’s multiple correction test. Different alphabetic letters show significant difference (P < 0.05) for each sex whereas, same alphabets denotes insignificance.

**S3 Fig. Effect of litter expansion on gene expression of IGF-1 levels in liver, gastrocnemius and brain cortex**. mRNA levels of IGF-1 in the liver (A), gastrocnemius (B) and brain cortex (C) of female and male mice from various litter sizes (LS6, LS8, LS10 and LS12 pups/litter) were measured at 15 months of age. Data represented are the mean ± SEM from 3-5 mice per group and were statistically analyzed by one-way ANOVA with Tukey’s multiple correction test. Different alphabetic letters show significant difference (P < 0.05) for each sex whereas, same alphabets denotes insignificance.

